# High-throughput detection of RNA processing in bacteria

**DOI:** 10.1101/073791

**Authors:** Erin E. Gill, Luisa S. Chan, Geoffrey L. Winsor, Neil Dobson, Raymond Lo, Shannan J. Ho Sui, Bhavjinder K. Dhillon, Patrick K. Taylor, Raunak Shrestha, Cory Spencer, Robert E. W. Hancock, Peter J. Unrau, Fiona S L Brinkman

## Abstract

**Background:** Understanding the RNA processing of an organism's transcriptome is an essential but challenging step in understanding its biology. Here we investigate with unprecedented detail the transcriptome of *Pseudomonas aeruginosa* PAO1, a medically important and innately multi-drug resistant bacterium. We systematically mapped RNA cleavage and dephosphorylation sites that result in 5'-monophosphate terminated RNA using a new high-throughput methodology called monophosphate RNA-Seq (pRNA-Seq). Transcriptional start sites (TSS) were also mapped using differential RNA-Seq (dRNA-Seq) and both datasets were compared to conventional RNA-Seq performed in a variety of growth conditions.

**Results:** The pRNA-Seq transcript library revealed known tRNA, rRNA and tmRNA processing sites, together with previously uncharacterized RNA cleavage events that were found disproportionately near the 5' ends of transcripts associated with basic bacterial functions such as oxidative phosphorylation and purine metabolism. The majority (97%) of the processed mRNAs were cleaved at precise codon positions within defined sequence motifs indicative of distinct endonucleolytic activities. The most abundant of these motifs corresponded closely to an *E. coli* RNase E site previously established *in vitro*. Using the dRNA-Seq library, we performed an operon analysis and predicted 3,159 potential TSS. A correlation analysis uncovered 105 antiparallel pairs of TSS that were separated by 18 bp from each other and that were centered on a palindromic TAT(A/T)ATA motif, suggesting that such sites may provide a novel form of transcriptional regulation. TSS and RNA-Seq analysis allowed us to confirm expression of small non-coding RNAs (ncRNAs), many of which are differentially expressed in swarming and biofilm formation conditions.

**Conclusions:** This study introduces pRNA-Seq methodology, which provides the first comprehensive, genome-wide survey of RNA processing in any organism. As a proof of concept, we have employed this technique to study the bacterium *Pseudomonas aeruginosa* and have discovered extensive transcript processing not previously appreciated. We have also gained novel insight into RNA maturation and turnover as well as a potential novel form of transcription regulation.

NOTE: All sequence data has been submitted to the NCBI short read archive. Accession numbers are as follows: [NCBI short read archive: SRX156386, SRX157659, SRX157660, SRX157661, SRX157683 and SRX158075]. The sequence data is viewable using Jbrowse on www.pseudomonas.com (example: http://tinyurl.com/pao1-prna-seq).

(An example of certain tracks is shown for convenience, but other tracks of data can be displayed using the “select tracks” option, and tracks may be clicked on and dragged to re-order them.)

## BACKGROUND

*Pseudomonas aeruginosa* is a medically important γ-proteobacterium that is noted for causing opportunistic infections in hospitalized patients and chronic lung infections in cystic fibrosis patients [1]. A substantial cause of human morbidity and mortality, *P. aeruginosa* has been broadly studied due to its metabolic diversity and its ability to undergo substantial lifestyle changes that include biofilm formation, swarming motility and quorum sensing, adaptive responses to antibiotics, and complex virulence adaptations [1]. While *P. aeruginosa* PAO1 is the type strain for this model organism, detailed knowledge of transcriptional start sites (TSS) is currently lacking for this isolate. The post transcriptional modifications of RNA transcripts are largely unknown in *Pseudomonas* and are generally poorly studied for any living organism. In addition to enhancing our understanding of the basic biology of *P. aeruginosa*, the detailed mapping of TSS and subsequent RNA processing of transcripts involved in virulence, antimicrobial resistance, and essential cellular functions, will aid in understanding the regulation of pathogenesis and drug resistance, and facilitate the identification of promising drug targets.

A systematic inventory of TSS and RNA processing sites is fundamental to understanding a broad range of cellular processes. Determining the set of post-transcriptional modifications found in the transcriptome of an organism is an important and yet very challenging objective that can be partially addressed by RNA sequence-based analysis. After transcription, a series of highly regulated secondary modifications occur that result in the maturation of an RNA transcript. These processing steps strongly influence the overall lifetime of the RNA molecule and are instrumental in the functionality of many RNAs [2, 3]. A primary transcript contains a terminal 5’ triphosphate [4] (Fig. 1). In bacteria, selective removal of the 5’ triphosphate by the conserved pyrophosphatase YgdP leaves a 5’ monophosphate [5]. This destabilizes mRNAs by making them more susceptible to 5’->3’ exonuclease degradation. An important complex in this regard is the multi-subunit degradosome, which contains the 5’ phosphate sensitive exonuclease/endonuclease RNase E at its core [3]. Similarly endonucleases that cleave RNA so as to leave a 5’ phosphate can either activate RNA for degradation via pathways that include the degradosome, or can result in the production of stable RNAs essential for cellular function. Transfer RNAs and the 4.5S RNA tmRNA that rescues ribosomes found on broken messages are all endolytically cleaved by RNase P, producing a stable 5’ phosphate terminus [3]. RNase E/G, which performs endolytic cleavage at A/U rich ssRNA regions, plays a central role in the 5’ maturation of the 5S RNA as well as the maturation of the 3’ ends of tRNAs [3]. Concurrently, RNase III serves to cleave dsRNA and has a primary role in the maturation of the 16S and 23S rRNAs [6]. The complex interplay between these and other endo- and exo-nucleases presumably acts on numerous other unstudied RNAs within a cell, helping to regulate the maturation and lifetime of expressed RNA. Studying such processing with high-throughput methodologies provides a significant window into understanding global aspects of transcriptional regulation.

**Figure 1:**
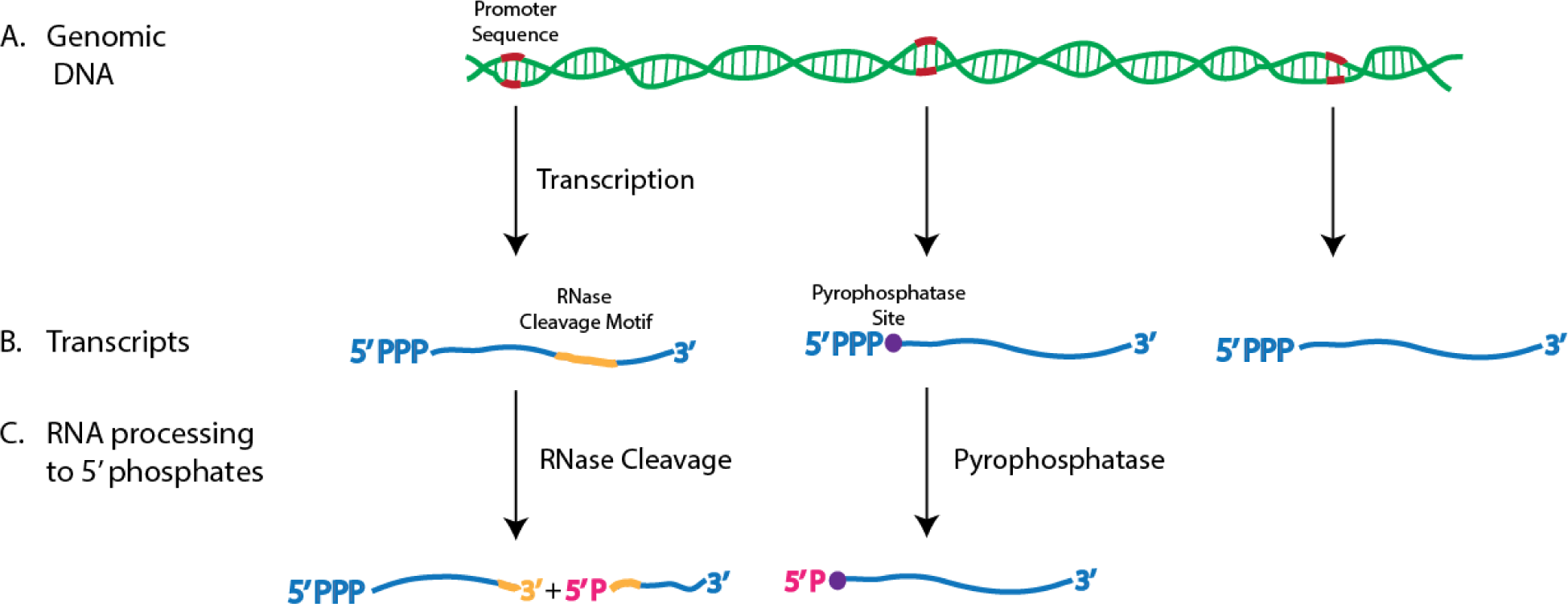
RNA transcription and processing. (A)Transcription of RNA is initiated from a promoter sequence (indicated in red) within the genome. Ribonucleoside triphosphate polymerisation results in a 5' triphosphate at the 5' end of the nascent transcript and a 3' hydroxyl at its 3' terminus. (B) mRNAs can be internally cleaved by endonucleases to yield two RNA fragments or can be degraded by exonucleases from either their 5' or 3' termini. The activity of exonucleases is often triggered by the selective dephosphorylation of a terminal triphosphate to a monophosphate by a pyrophosphatase. (C) This study focuses on all RNA processing event that result in either a 5' triphosphate (dRNASeq) or 5' monophosphate (pRNA-Seq) and that simultaneously contain a terminal 3' hydroxyl.

RNA transcription is one of the initial steps in a complex regulatory cascade that enables cells to synthesize and regulate the expression of cellular factors in response to environmental changes. The highly conserved process of sigma factor dependent transcriptional initiation is of central importance in all bacteria [7]. Sigma factors form part of the RNA polymerase holoenzyme during transcriptional initiation and determine which promoters are active in specific cellular states [8]. *P. aeruginosa* PAO1 has 24 putative sigma factors, of which 14 have yet to have their DNA binding sites identified [9]. These sigma factors and their associated regulators are responsible for the correct transcriptional response to changing environmental conditions including low oxygen [10], limited iron [11, 12], and overall nutrient levels [13]. Identifying TSS within the genome and defining upstream sequence motifs helps to identify sigma-dependent promoters. To date, only 83 TSS have been annotated in the PAO1 strain [14, 15]. Wurtzel et al. [16] performed a key expansion of annotated TSS in *P. aeruginosa* strain PA14 by employing dRNA-Seq to find 2,117 putative TSS. Notably however, the PA14 strain differs from the PAO1 strain [17] in that it contains an additional ∼200 genes, is known to be more virulent [18], and has an estimated 5,977 open reading frames versus the 5,688 in PAO1 [1]. There is therefore considerable benefit in systematically defining and exploring TSS in PAO1.

The dRNA-Seq method was pioneered [19] to comprehensively map TSS found in a prokaryotic genome that are expressed in a given condition. This was accomplished by sequencing, in an orientation specific manner, RNA transcripts with triphosphates (PPP) at their 5’ ends. RNA-Seq technology has quickly become the new standard in transcriptome analysis and the data derived from these experiments has allowed us to view transcriptomes in unparalleled detail (see, e.g. [16, 20–23]). Single base pair resolution maps of transcriptional products derived from high throughput sequence data allow for gene by gene quantification of expression levels and novel gene discovery. However, 5’ degradation occurs quickly in bacterial RNA samples and it is difficult to tell where TSS are located based on standard RNA-Seq data. dRNA-Seq allows for the identification of TSS by sequencing only those transcripts that contain triphosphates at their 5’ ends [19], but this method cannot be used to examine further processing of transcripts after their synthesis by RNA polymerase. Thus our current picture of RNA processing in bacteria is substantially incomplete.

We report here a novel methodology, monophosphate RNA-Seq (pRNA-Seq), and use it to study RNA processing in *P. aeruginosa* strain PAO1. In addition, we have used the dRNA-Seq methodology of Sharma et al. [19] to characterize TSS and have conducted RNA-Seq inventories under four different growth conditions, in addition to selected additional downstream experiments, to provide a more complete picture of RNA expression in this organism. In addition to locating 1,741 5’ monophosphate cleavage sites, we have also identified the sequence motifs corresponding to these sites, and were able to propose specific nucleases that might be responsible for some of the observed cleavage events. In addition we identified 3,159 probable TSS in PAO1, significantly expanding our understanding of TSS in *P. aeruginosa*. A fraction of these TSS was found to be arranged in antiparallel pairs, implying that transcriptional initiation at either site might be conditionally dependent on the other. Through further downstream experiments, we demonstrated that certain small non-coding RNAs (ncRNAs) show significant changes in expression during swarming and biofilm formation, suggesting important roles for these RNAs in determining these complex adaptations. Collectively, these studies have heightened our understanding of transcription and RNA processing in the s-proteobacteria, revealing layers of RNA processing complexity that were previously unexplored.

## RESULTS

Our transcriptome analysis consisted of three main facets: First, we sequenced the genome of our isolate of PAO1-UW using the Illumina methodology to confirm that it is the ref-seq isolate [1]. Our PAO1-UW isolate was nearly identical to the published reference sequence NC_002516, differing by only 11 single nucleotide polymorphisms (SNPs), each of which had been previously shown to be variable among laboratory strains of PAO1-UW [24], as well as 10 small indels (See Tables S1A and S1B respectively for a summary). The reference sequence genome was then used as a basis to map all reads from libraries derived from the transcriptome of PAO1-UW. Second, we analyzed the transcriptome for RNAs terminated with a 5’ monophosphate using our new high throughput method, pRNA-Seq. Third, we used dRNA-Seq to identify TSS and supplemented this information with an RNA-Seq analysis of transcription under four distinct growth conditions selected to simulate both laboratory and a range of infectious conditions. For all libraries, reads were then mapped to either the plus or minus strands (except for the RNA-Seq data, where information on strand orientation was not available) of the PAO1-UW genome using criteria summarized in the methods. We integrated data from the 5’ termini of the strand orientation sensitive dRNA-Seq and pRNA-Seq libraries into a detailed map of the PAO1-UW genome that identified both cleavage sites and TSS. RNA-Seq data was used to calculate read depth (see Methods) that again typically correlated well with both TSS and sites of major RNA processing as determined by dRNA-Seq and pRNA-Seq respectively. Statistics on the size and composition of each library can be viewed in Table 1. For online Jbrowse access to our data sets see www.pseudomonas.com

**Table 1:** Library growth conditions and summary of the number and percentage of reads mapped to the *Pseudomonas aeruginosa* PAO1 reference sequence and including (+ RNA) or excluding (- RNA) rRNA, tRNA and tmRNA genes. In all libraries but 110817_SN865, which consists of single end reads, the mapped reads are in proper pairs with a maximum insert size < 1,000.

| Sample | Total Reads | including rRNA |  | excluding rRNA |  |
| --- | --- | --- | --- | --- | --- |
|  |  | Reads Mapped | Percent Mapped | Reads Mapped | Percent Mapped |
| pRNA-Seq<br>LB Medium, 37°C, OD<sub>600</sub> = 0.7<br>(A06027) | 272,983,632 | 149,142,710 | 55% | 51,067,158 | 19% |
| dRNA-Seq<br>LB Medium, 37°C, OD<sub>600</sub> = 0.7<br>(110817_SN865) | 131,130,793 | 108,900,693 | 83% | 29,801,108 | 23% |
| RNA-Seq<br>Synthetic Cystic Fibrosis Medium,<br>37°C, OD<sub>600</sub> = 0.7 (A03674) | 54,000,716 | 51,910,678 | 96% | 16,003,536 | 30% |
| RNA-Seq<br>Artificial Sputum Medium, 37°C, OD<sub>600</sub><br>not determined (A06026) | 63,854,706 | 61,850,374 | 97% | 6,802,731 | 11% |
| RNA-Seq<br>LB Medium, 37°C, OD<sub>600</sub> = 0.7<br>(PA0004) | 75,437,354 | 72,729,310 | 96% | 30,068,678 | 40% |
| RNA-Seq<br>LB Medium, 34°C, OD<sub>600</sub> = 0.7<br>(PA0001) | 20,643,394 | 17,559,382 | 85% | 3,352,489 | 16% |

### Terminal monophosphate RNA data analyzed by pRNA-Seq revealed both expected and novel transcriptome processing sites

RNA processing sites are defined here as genomic locations with R100 reads “first bp coverage” from the pRNA-Seq library (i.e. oriented and aligned sequences whose 5’ most nucleotide was found at least 100 times; such sequences can and often had non-homogenous 3’ ends–see Methods). Processing sites were found that corresponded to previously-characterized processing sites of RNAs involved in translation. The *ssrA* RNA (tmRNA) was matured at precisely the residue expected based on data from other s-proteobacteria (Fig. 2A) [25], as were the 12 ribosomal RNAs (4 copies of each of 5S, 12S and 23S RNA) [6]. However, the ribosomal RNAs also contained multiple internal cleavage sites. This was unexpected, as ribosomes are believed to be relatively stable structures within the cell, with core rRNAs that are resistant to the action of nucleases. The degradation process of ribosomes has, however, not been intensively studied [6], and it is currently unclear exactly what fraction of ribosomes are degrading during normal growth conditions. We examined the forty-one tRNAs known to be expressed in the pRNA-Seq library growth conditions (from our LB 37^0^C RNA-Seq data) for cleavage sites. Isoleucine, alanine, serine and leucine tRNAs were observed to be processed to monophosphates at their 5’ termini, as would be expected based on the activity of RNase P. Overall a total of 13 tRNAs were cleaved at some point within the transcript, and by lowering the cut off threshold for cleavage site prediction (see Methods) from 100 reads to 50 reads, 5 additional tRNAs were revealed to be subject to cleavage. Transfer RNAs within an operon containing multiple tRNAs (i.e. Fig. 2B) showed evidence of not only 5’ processing but also 3’ processing and occasionally internal tRNA cleavage; this has been previously described as a mode of tRNA regulation in other bacteria, as well as in eukaryotes [26, 27]. Typically tRNA processing signals were more prominent towards the 5’ ends of the operons as would be expected from our use of a random primer reverse transcription (RT) step to generate cDNA after the initial ligation of a 5’ adapter sequence. Given that our RNA isolation method preferentially excluded tRNAs, it is unsurprising that they were not always present in large enough quantities to pass the step size threshold filtering steps for the pRNA-Seq library. Together, these data served as important biological controls for the overall pRNA-Seq methodology.

**Figure 2:**
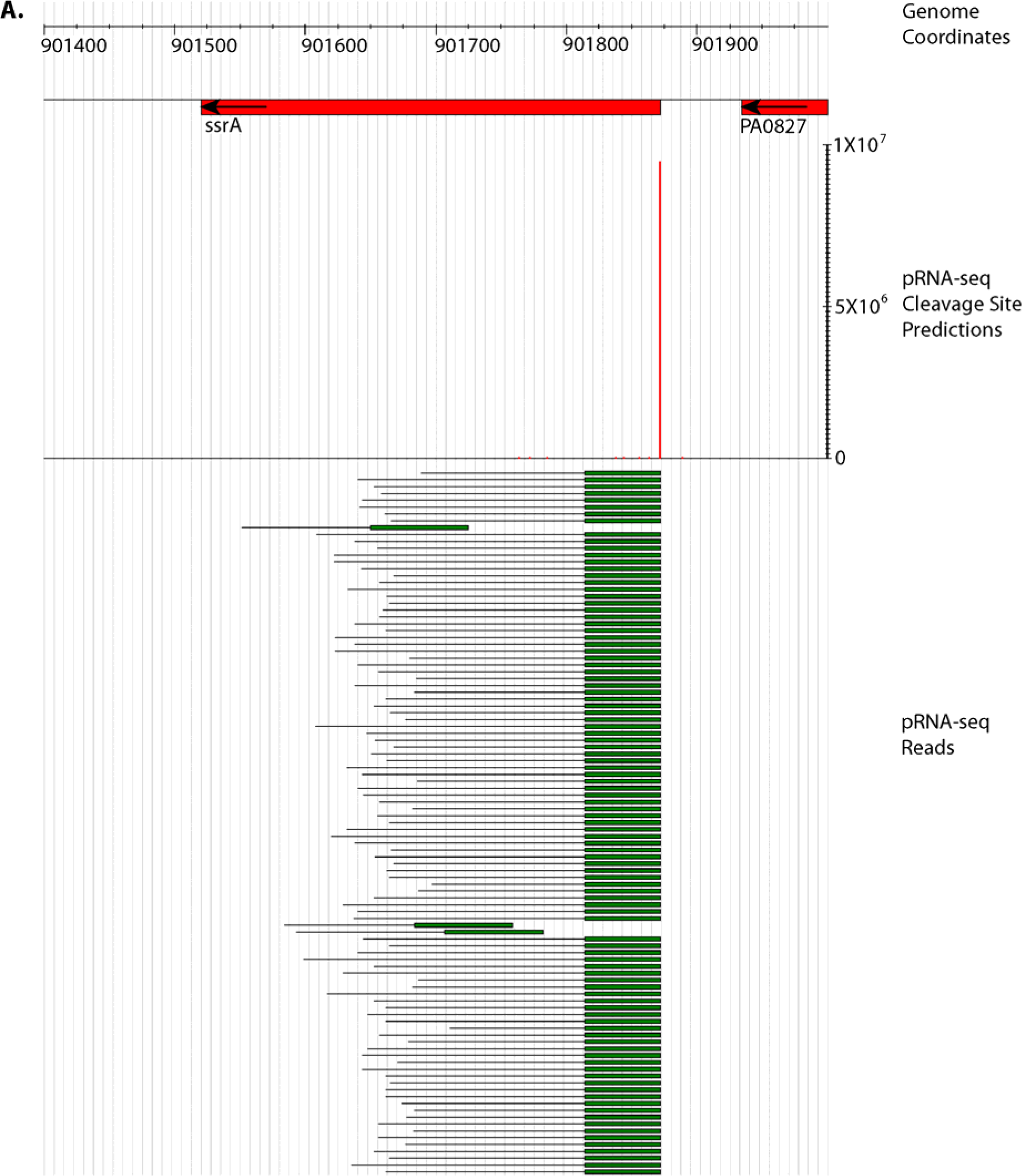
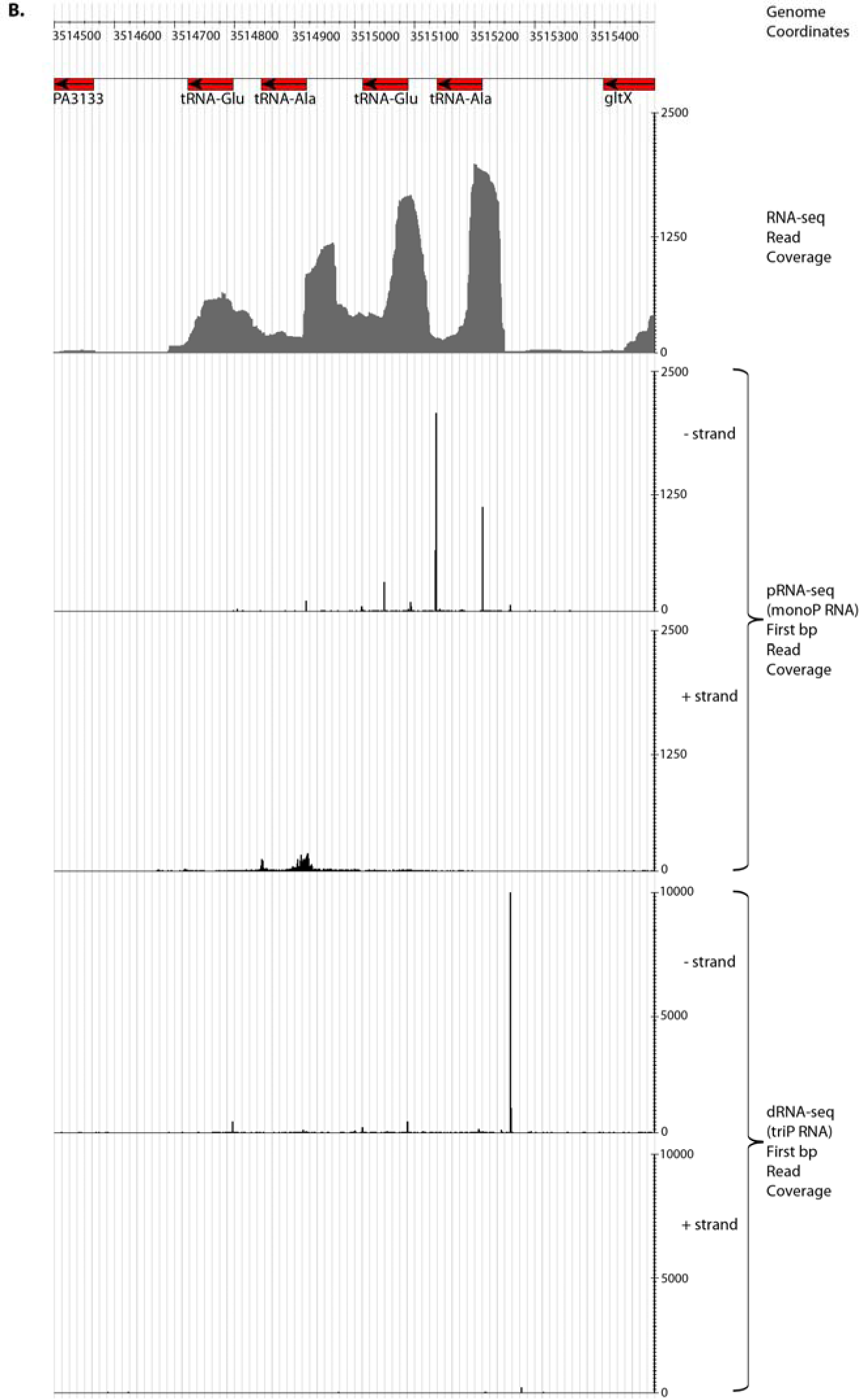
Identification of precise RNA processing events within the ssrA transcript (the tmRNA) and tRNA operon transcripts. (A) A histogram showing precise processing of the ssrA gene (by RNase P), with the 5’ ends of processed pRNA-Seq paired end reads aligning exactly at a single genomic location. Transcription for this gene is initiated 60-nt upstream of the location of RNase P cleavage. (B) Conventional RNA-Seq suggests the possibility that complex RNA processing occurs within the tRNA operon shown. pRNA-Seq indicates that a series of precise RNA cleavages occur downstream from a single strong initial transcriptional start.

In addition to finding the expected evidence of RNA processing in certain transcripts, we observed evidence that unique cleavage events occurred in a diverse set of transcripts. Overall we identified 1,741 5’ monophosphate RNA processing sites in the PAO1 transcriptome that met our statistical criteria for significance (Fig. 3). 500 putative cleavage sites not overlapping with pyrophosphatase sites were identified in 240 protein-coding mRNAs, which were among the most heavily transcribed in both the dRNA-Seq and RNA-Seq LB 37^0^C libraries. Of the 240 most abundantly transcribed protein coding genes in the LB 37^0^C RNA-Seq library (Table S2), transcripts from 111 of these genes, or 46%, were also present in the dRNA-Seq library. Forty three (nearly 18%) of these genes were ribosomal proteins, which was by far the most highly represented Kyoto Encyclopedia of Genes and Genomes (KEGG; [28]) category among the transcripts. Three ncRNAs (PA4406.1, *rnpB* and *crcZ*) also showed evidence of cleavage. In addition, we found clear evidence of cleavage in transcripts for a wide range of proteins associated with critical cellular functions other than protein translation. For example, cleavage in PA3648/ *opr86* was detected, particularly in the central region of the gene. PA3648 encodes the only essential integral outer membrane protein in *P. aeruginosa* PAO1–a protein critical for outer membrane biogenesis [29].

**Figure 3:**
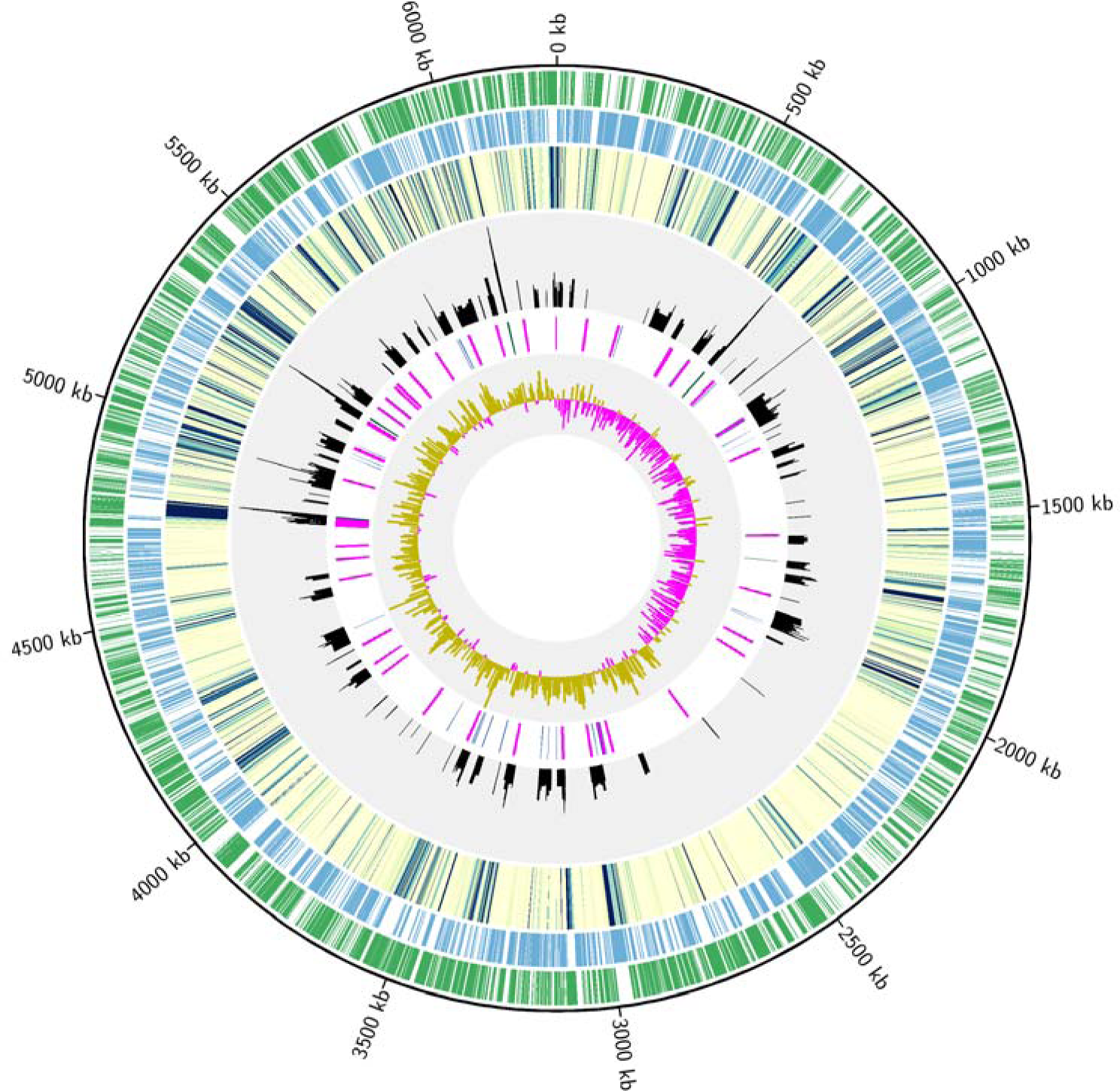
Circular plot showing distribution of mapped RNA-Seq reads and 5’ monophosphate cleavage sites throughout the *Pseudomonas aeruginosa* PAO1 genome. The outer green and blue tracks represent reverse-strand and forward-strand genes, respectively. The third track from the outside is a heat map showing log10 first base-pair coverage (100 bp window) of RNA-Seq reads where a transition from yellow to green to blue correlates with increased transcription. The histogram with a grey background shows log10 coverage of 5’ monophosphate sites on both the forward and reverse strands. This is followed by a track containing rRNA (green), tRNA (blue) and ribosomal proteins (purple). The innermost track shows G-C skew (1,000 bp window).

### RNA cleavage sites within annotated genes were often clustered in the 5' untranslated region of transcripts and were correlated with the reading frame position

Figure 3 presents a global overview of the locations of cleavage sites within the genome. The RNA cleavage site locations within protein-coding genes were analyzed relative to the start and stop sites of annotated protein coding genes. This revealed that 31.6% (158/500) of cleavage sites fell upstream of the translation initiation site, with the remaining cleavage sites being nearly uniformly distributed across genes (Figure 4A). We sought to determine whether there was a correlation between ribosomal binding site (RBS) location and cleavage site location. Prodigal software [30] was used to predict RBSs upstream of PAO1 protein coding sequences. Of the genes possessing a recognizable RBS and for which we had both TSS and RNA cleavage data, 2% (11) produced transcripts that were cleaved either within or immediately downstream of (i.e. 3’ to) the RBS. The ORFs of such cleaved RNAs would presumably lack a RBS and consequently be expected to be poorly translated. More than half of the cleavage sites within gene ORFs were associated with a particular codon position, with 267 out of 500 (53.4%) of the ORF cleavage sites being located immediately following the first base in a codon, 5’-N1^N2N3 (^: site of cleavage, Fig. 4B). This is likely due to the notable G+C bias in the genome influencing the location of G and C nucleotides in the cleavage site motif. Most of the cleavage sites located within genes were disproportionately located within transcripts for genes associated with specific functional categories as defined by the KEGG [28]. These included the basic bacterial functions of oxidative phosphorylation (54 cleavage sites, 14 genes, corrected p-value =0.0000049) and purine metabolism (37 cleavage sites, 13 genes, corrected p-value = 0.0021), indicating a potentially important role in the regulation of core metabolism by the post-transcriptional secondary cleavage of RNA transcripts. Protein coding genes that contained cleavage sites tended to code for essential cellular functions. These genes encoded proteins that acted the toluene degradation pathway (3 genes, corrected p-value 0.004.4), and RNA polymerization (2 genes, corrected p-value 0.015).

**Figure 4:**
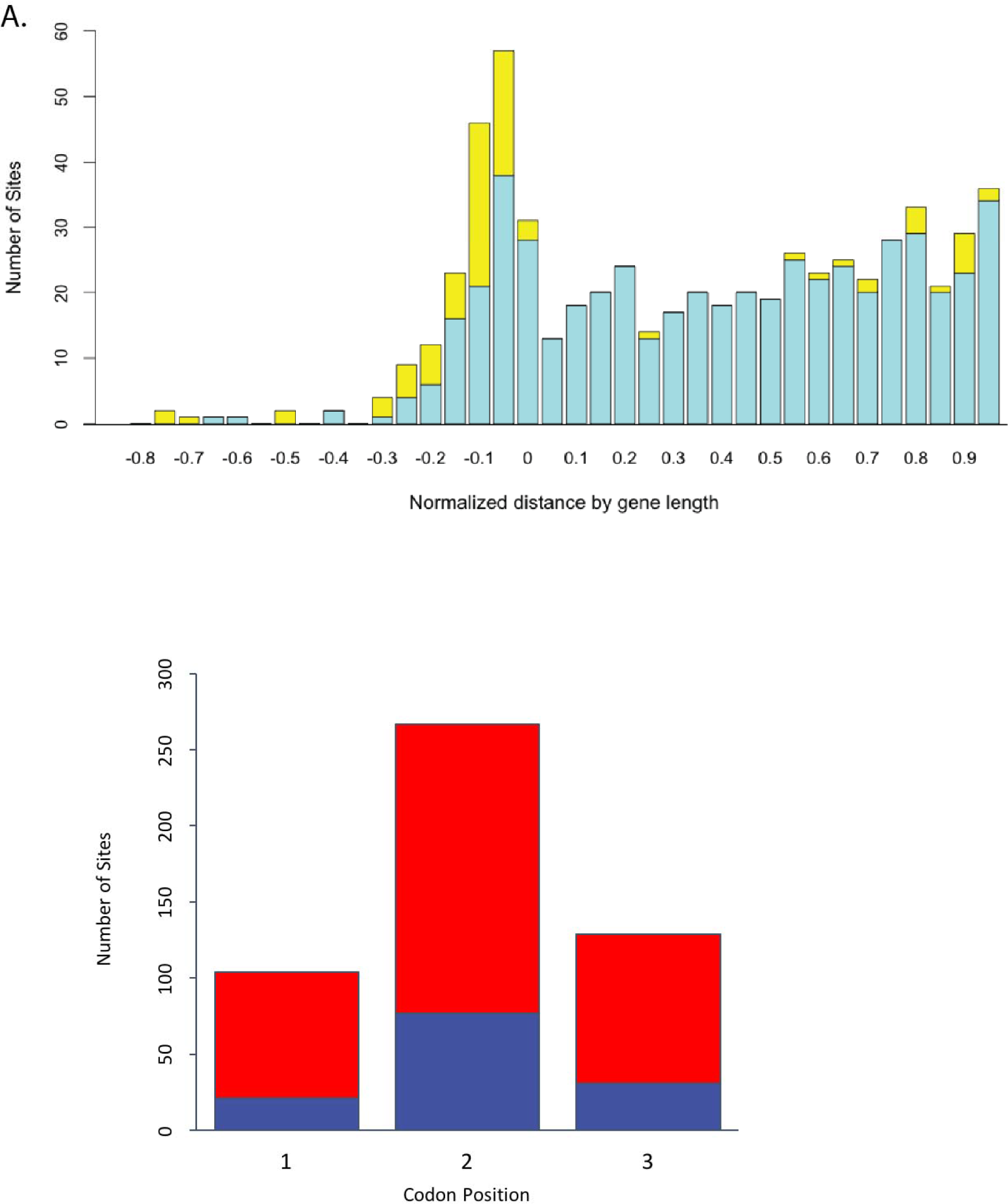
Relative cleavage site position within genes and reading frame dependent cleavage bias. (A) The relative distance that cleavage sites fall within protein coding genes was plotted in a histogram (see materials and methods). Cleavage sites are shown in turquoise, 5' monophosphate sites occurring at the location of a TSS (i.e. potential 5' pyrophosphatase sites) are shown in yellow. The remaining sites are distributed relatively evenly throughout the ORF.(B) The reading frames of cleavage products were determined for both positive (navy blue) and negative (red) stranded transcripts. Cleavage at site 1 occurs 5’ to nucleotide 1 ( N1N2N3), cleavage at site 2 occurs 5’ to nucleotide 2 (5’-N1-N2N3) and cleavage at site 3 occurs 5’ to nucleotide 3 ( 5’-N1N2-N3).

### RNA cleavage patterns correlated with RNA cleavage motifs

We next searched for RNA sequence motifs associated with cleavage sites within annotated genes to determine whether specific nucleases might be involved in their creation. Processing sites found at locations meeting our peak height cut off criteria, and not overlapping with pyrophosphatase sites were aligned and examined for potential patterns associated with nuclease digestion. These potential patterns were inspected in a 10-nt window upstream and downstream of the dominant cleavage site which had the highest read coverage in the window. This alignment was used to explore the hypothesis that distinct RNA digestion patterns might be correlated with specific RNA sequence motifs. Strikingly, a single global motif dominated the dataset (Fig. 5A), [(A,C,g,u)(A,C,g,u)(G,a,c)(A,g,u)↓(A,c,u)(**C**,u)(A,c,g)(C,a,g,u)(C,a,g)], wherein nucleotides present in 10-30% of the sequences are depicted in lower case letters, nucleotides present in 31-64% of the sequences are depicted in uppercase letters and nucleotides present in 65% or more of the sequences are depicted in bold upper case letters. The position of the predominant cleavage site is indicated by the downward arrow (see also the graphical views of motifs in Figures 5 and S1).

**Figure 5:**
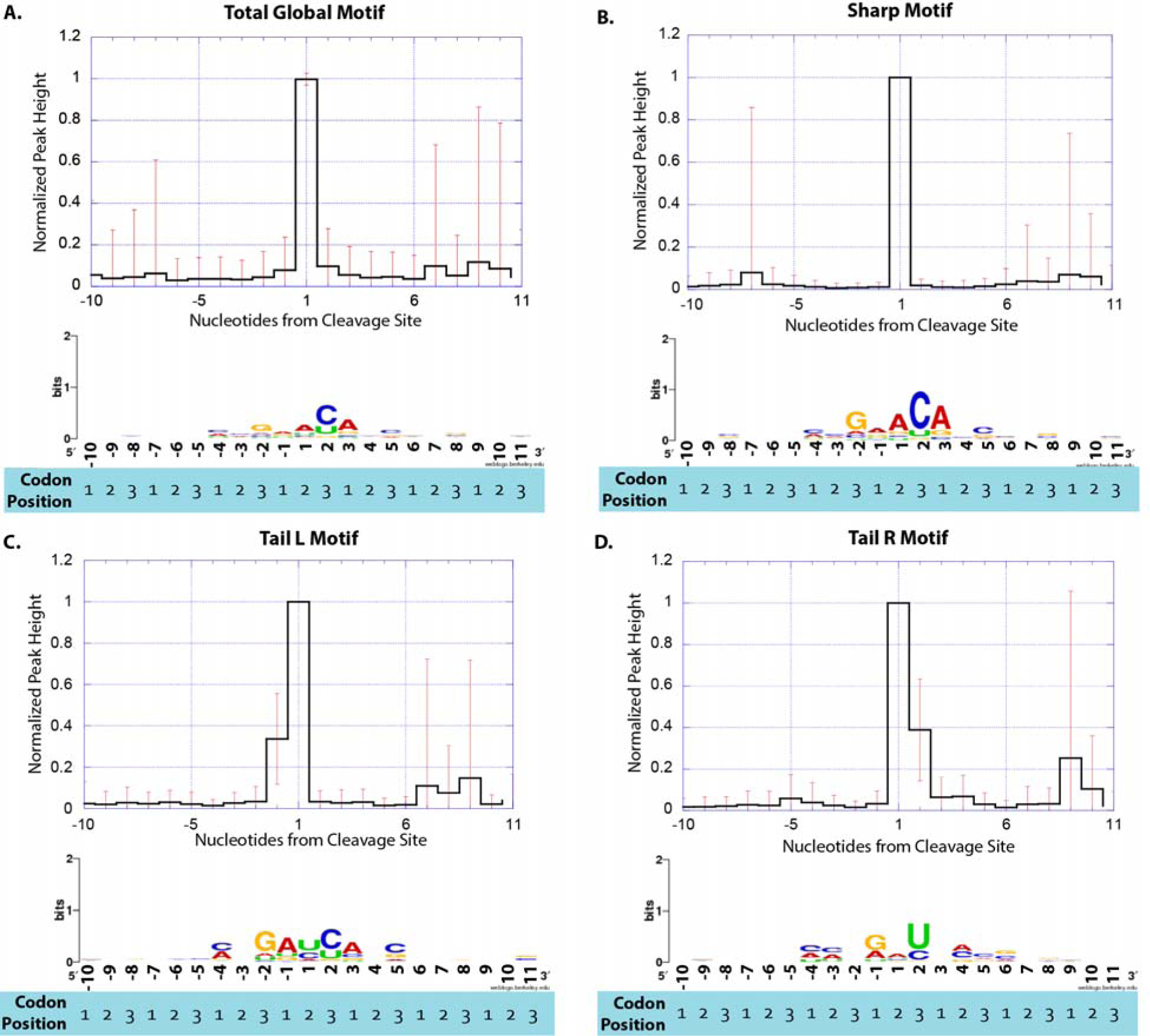
5’ Monophosphate processing patterns and their corresponding sequence motifs. Cleavage sites are derived from genome locations with 100 reads or more 1^st^ bp coverage from our pRNA-Seq library. Peak shape refers to the number of mapped transcripts surrounding a cleavage site. Peak shapes were categorized using k-mean clustering. Motifs were calculated from peak shape clusters using MEME [58]. The graph at the top of each panel shows normalized peak height. The WebLogo [59] at the bottom of each panel shows the sequence motif associated with each peak shape. (A) The global motif, which is derived from the entire pRNA-Seq dataset. (B) the “Sharp” peak shape motif, shows strong similarity to the RNase E motif in *E. coli*[36]. (C) the “Tail L” motif and (D) the “Tail R” motif.

K-means clustering decomposed the RNA digestion patterns into five distinct classes based on peak shape that correlated with specific RNA sequence motifs. By peak shape we mean the shape of the pattern produced by the mapped 1^st^ bp coverage of transcripts surrounding a cleavage site. For more in-depth analysis of motifs surrounding cleavage sites, we chose to analyze cleavage sites that fall within ORFS and are >= 10 nt downstream of another peak. Of the 383 cleavage sites analyzed, 97% fell into one of the categories described below. For a list of all RNA sequence cleavage motifs, see Table S3. The predominant ‘Sharp’ RNA cleavage pattern (182/383 cleavage sites) consisted of a single RNA cleavage event with little or no cleavage at adjacent nucleotides, and therefore resembles a sharp peak when viewed graphically (Fig. 5B). The Sharp sequence motif was consistent with the overall global motif, containing the nucleotides [(A,C,g,u)(A,C,u)(**G**,a,c)(A,g,u)↓(**A**,c,g)(**C**,u)(**A**,c,g)(C,a,g,u)(C,a,g)], where the downward arrow indicates the cleavage site. RNAs that contained this cleavage motif were located in genes that disproportionately belonged to the KEGG categories “oxidative phosphorylation” (number of genes = 12, number of cleavage sites = 54, corrected p-value =0.000026) and “purine metabolism” (number of genes = 12, number of cleavage sites = 37, corrected p-value = 0.0011) (Table 2).

**Table 2:** Association of cleavage site motif types with KEGG functional terms. Certain cleavage site motifs are disproportionately located within genes that are associated with certain KEGG functional terms. The following table lists the KEGG terms associated with each cleavage site motif type and the p-values associated with each KEGG term. P-values were calculated from hypergeometric tests for KEGG pathways and Holm’s test was used to correct for multiple testing.

| **Test by peak shape** | **KEGG functional terms** | **p value** | **Number of genes in set with this function** | **Number of Cleavage Sites by KEGG Term** |
| --- | --- | --- | --- | --- |
| Global | Oxidative phosphorylation | 4.90E-06 | 14 | 54 |
|  | Purine metabolism | 2.08E-03 | 13 | 37 |
|  | Aminoacyl-tRNA biosynthesis | 8.94E-03 | 7 | 16 |
|  | RNA polymerase | 1.72E-02 | 3 | 19 |
|  | Pyrimidine metabolism | 4.67E-02 | 8 | 26 |
|  | Protein export | 4.93E-02 | 5 | 37 |
| Sharp | Oxidative phosphorylation | 2.60E-05 | 12 | 54 |
|  | Purine metabolism | 1.13E-03 | 12 | 37 |
|  | RNA polymerase | 9.13E-03 | 3 | 19 |
|  | Pyrimidine metabolism | 1.14E-02 | 8 | 26 |
|  | Aminoacyl-tRNA biosynthesis | 1.89E-02 | 6 | 16 |
| Tail L | RNA polymerase | 7.11E-05 | 3 | 19 |
|  | Protein export | 1.24E-02 | 3 | 37 |
|  | Purine metabolism | 1.39E-02 | 5 | 37 |
| Tail R | RNA polymerase | 4.52E-02 | 3 | 54 |
|  | Protein export | 1.51E-02 | 2 | 37 |
| Twin R | Oxidative phosphorylation | 9.70E-04 | 5 | 54 |
|  | Protein export | 1.03E-02 | 3 | 37 |
|  | Oxidative phosphorylation | 2.52E-02 | 4 | 54 |
|  | Citrate cycle (TCA cycle) | 4.94E-02 | 3 | 39 |

The remaining cleaved RNAs sorted into RNA digestion patterns that showed asymmetries in their cleavage patterns (Figs. 5C and D). When viewed graphically, the second most abundant peak shape (58/383 cleavage sites) had a shoulder immediately 5’ to the dominant peak (Fig. 5C) and was named “Tail L”. This motif (Fig. 5C) contained the nucleotides [(A,C)(A,c,g,u)(**G**,a,u)(A,g,u)↓(C,U,a)(C,u)(A,c,g)(A,C,g,u)(C,a,g)] and was consistent with either two adjacent cut sites or an initial cleavage event followed by the removal of an additional nucleotide in the 5’->3’ direction. The motif for this cluster shared properties with the predominant Sharp motif but was notably lacking in sequence conservation at the +2 and +3 position. The Tail L motif also frequently (>30%) had a U at the +1 position that was absent in the predominant Sharp motif. Transcripts containing this cleavage motif included those from genes belonging to the KEGG categories “RNA polymerase” (number of genes = 3, number of cleavage sites = 19, corrected p-value = 0.000071), “protein export” (number of genes = 3, number of cleavage sites = 37, corrected p-value = 0.012) and “purine metabolism” (number of genes = 5, number of cleavage sites = 37, corrected p-value = 0.014).

The third most abundant peak shape (54/383 cleavage sites) had a shoulder immediately 3’ of its main peak and was named “Tail R” (Fig. 5D). This motif contained the nucleotides [(A,C,u)(A,C,u)(G,a,c,u)(A,G)↓(A,c,g,u)(**U**,c)(C,a,g,u)(A,C,g)(C,a,g)] and was quite different from the Sharp and Tail L motifs (Fig. 5C). RNAs containing this motif included transcripts from genes belonging to the KEGG categories “RNA polymerase” (number of genes = 2, number of cleavage sites = 19, corrected p-value = 0.0096) and “protein export” (number of genes = 3, number of cleavage sites = 37, corrected p-value = 0.010) among others (Table S3, Additional RNA cleavage patterns were also identified–see Supplementary Information for details).

A subset of the pRNA-Seq-determined cleavage sites were found to localize exactly with our determined TSS locations. In total 131 sites (92 in coding genes + 39 in rRNA) met this criterion and were thus likely to be due to dephosphorylation of transcripts whereby the triphosphate was removed from the 5’ end of the RNA molecule and a 5’ monophosphate remained (Fig. 1). One of the steps during preparation of the dRNA-Seq TSS libraries was the removal of 5’ monophosphate-containing RNA using terminator-5′-phosphate-dependent exonuclease. It was possible that some TSS might have been identified as false positives due to the incomplete digestion of 5’ monophosphate RNA during TSS library construction. However,if indeed removal of 5’ monophosphate RNAs was only partially complete, then we would have expected that the 5’ termini of tRNA and tmRNA, which are known to possess 5’ monophosphates, would be identified in our TSS (dRNA-Seq) library data. Since this was not observed, we can tentatively conclude that either the desphosphorylation sites identified were biologically significant, or that the secondary structures with the 5' termini of these RNAs might prevent dephosphorylation. Genes with transcripts that possessed dephosphorylation sites were not significantly associated with any KEGG terms. Due to the low number of dephosphorylation sites identified, motif analysis of the downstream sequence was inconclusive, as was an attempt to determine potentially conserved RNA secondary structure at these sites.

### TSS prediction from dRNA-Seq identified promoter regions and novel sets of potentially co-expressed genes

We predicted a total of 3,159 TSS from which RNA was actively initiated at 37^0^C in LB media. These sites were associated with 2,030 genes (in many cases, more than one TSS was associated with a single gene) that represented 36% of the strain PAO1 genome. Strikingly, just 54% (1,695) of these TSS lay outside of ORFs. The remaining 1,467 TSS lay within ORFs implying a potential regulatory role for such transcripts. We compared our predicted TSS to a set of 51 previously described TSS from strain PAO1 grown under various conditions [14], and for these genes 44 (86%) of our dRNA-Seq TSS lay within ±3 nt of previously published TSS (see Fig. S3).

### TSS correlations between plus and minus strands

A correlation analysis for TSS was performed to explore the hypothesis that adjacent TSS might be related to a particular biological process. This analysis revealed a novel and previously unsuspected correlation between back to back TSS pairs. These antiparallel TSS pairs, based on opposite strands, were separated by 18 bp (Figure 6A) and often contained a palindromic “TAT(T/A)ATA” motif equidistant between the two antiparallel TSS (Figure 6B). When a similar correlation analysis was performed for pairs of TSS on the same strand, no significant correlation between peaks was found (Figure 6A). A total of 105 antiparallel transcriptional pairs were found, with 59 of these sites being found between genes that were also in an antiparallel arrangement. Interestingly, 14 of these sites would be predicted to express antisense RNA for an otherwise parallel arrangement of genes.

**Figure 6:**
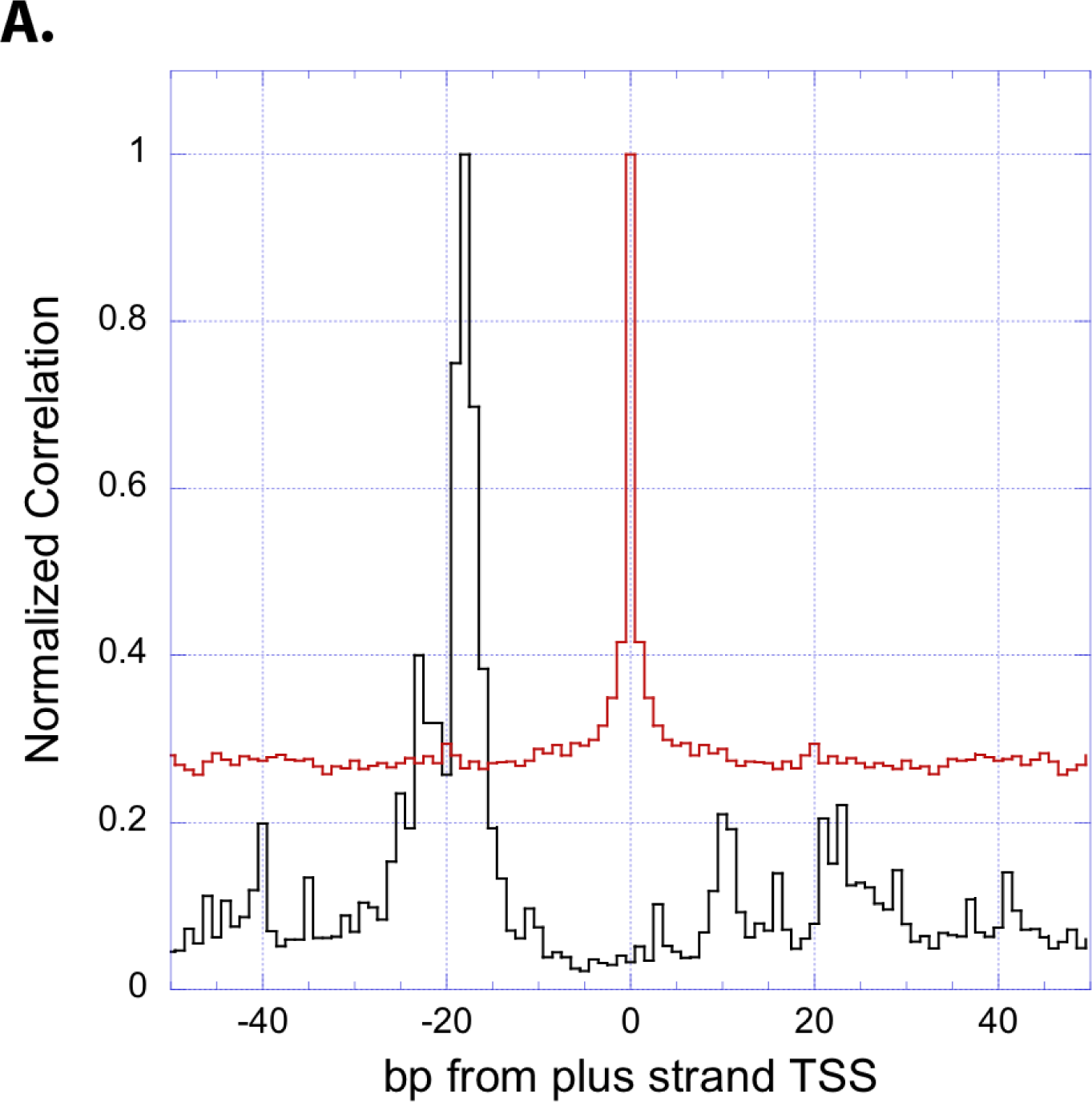
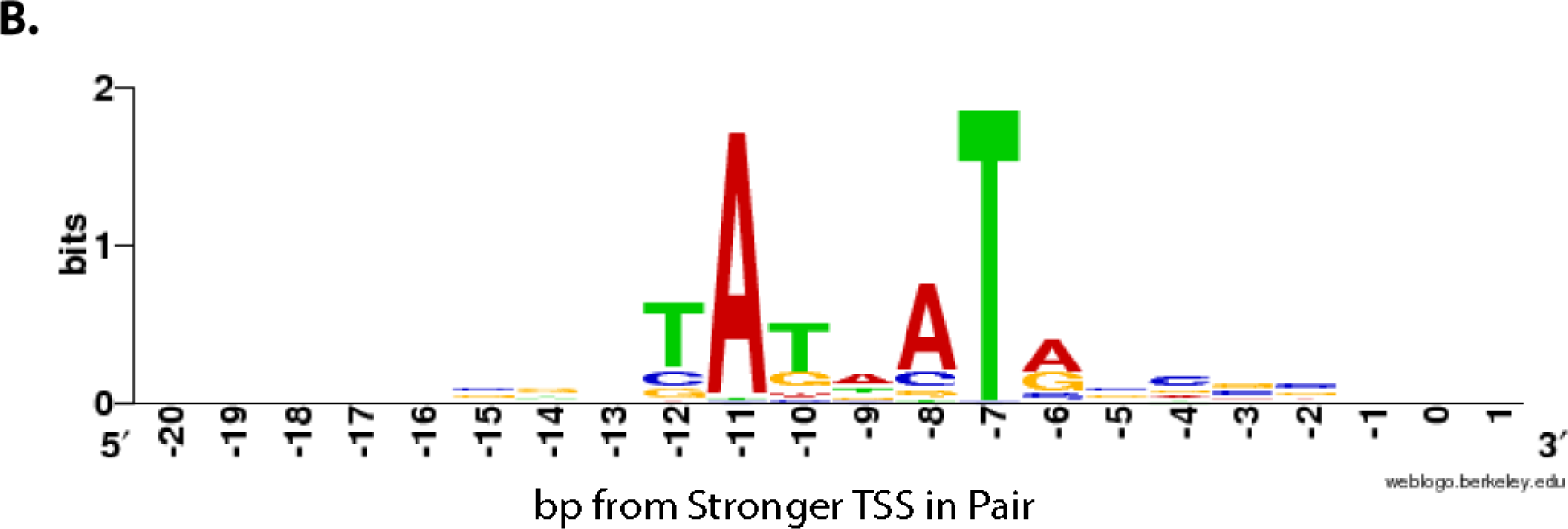
Correlation between TSS occurring in the antiparallel orientation and a palindromic sequence motif. (A) TSS show a significant correlation with TSS in the opposite direction that are 18-bp upstream (black curve, n=-18). No statistically significant correlation was found between TSS having the same strand orientation (red curve). The correlation function defined as: C(n) = Sum[x(l)y(l+n)], l=1..G-n, with G being the genome size, x(l) plus strand TSS counts, y(l) either minus strand TSS counts (black curve) or plus strand TSS counts (red curve). The maximum value of the correlation function was normalized to one for each curve. (B) A “TATNATA” motif occurs between antiparallel TSS. Of the 105 antiparallel TSS found that where 18-bp appart, 3 match the motif exactly, 27 have one mismatch, 25 have two mismatches and 13 have three mismatches.

### PAO1 transcriptional promoter map

The dRNA-Seq data analyzed here was used to develop a promoter map for strain PAO1. Promoters were predicted for all (1,612) primary TSS (TSS in an intergenic region and on same strand as downstream gene), including 111 primary antisense TSS (TSS in an intergenic region but on the opposite strand of the closest gene). Promoters can be viewed on Jbrowse at www.pseudomonas.com. Analyses were conducted to identify novel virulence factors based on the similarity of promoter motifs to known virulence factors. Three potential novel virulence factors that share promoter motifs with the gene *flhA* were identified (See Supplementary Material, Figure S4). In addition, we aimed to determine whether novel binding sites for the known sigma factor RpoN could be identified based on sequence motif similarity. 32 putative novel RpoN binding sites were identified within promoter regions of known genes. Four of these genes were predicted to be regulated by RpoN, but binding sites had not been described, while 25 were upstream of genes where RpoN involvement in transcription had not previously been hypothesized (See Supplementary Material).

### Chromosomal gene position affects transcription

Recently, it was reported that for short-read Illumina sequencing of bacterial genomes, sequence reads near the chromosomal origin of replication are more frequent than sequence reads distal to the origin. This is thought to be due to the nature of circular chromosome replication, such that there is a higher copy number of genes/sequences near the origin where DNA replication is initiated. Such read frequencies can even aid in the identification of genome rearrangements [31]. We examined whether RNA-Seq sequencing reads would have a corresponding bias towards higher frequency around the origin vs. the terminus of replication. RNA-Seq sequence reads did indeed show a decrease in frequency in a region near the known terminus (Fig. 3) that had been previously shown to also have reduced transposon mutagenesis frequency [32]. The fold-change of read density was also calculated in 0.5 Mbp increments along the genome when compared to the region with the lowest read density (Table S6). In all RNASeq libraries, the region with the lowest read density lay between 2 and 2.5 Mbp, which is the location of the terminus of replication as revealed by a G-C skew plot (Fig. 3). There was an array of rRNA genes located between 5 and 5.5 Mbp, which raised the fold-change of read density values to very high levels. In addition, the highly transcribed tmRNA gene was located at 1.1 Mbp, thus increasing fold-change values in this region. The regions with the highest read density were proximal to the origin of replication, having implications for the analysis of gene expression and illustrating the importance of gene location in impacting its expression.

### Confirmation, and additional functional analysis, of small non-coding (nc) RNAs

Estimates of the number of small, non-coding RNAs (ncRNAs) vary widely and the small size of ncRNA genes can make TSS mapping challenging. Gomez-Lozano et al. [33] utilized an automated analysis to identify 513 novel ncRNA transcripts and verified the expression of several previously-known ncRNAs. Another study by Wurtzel et al. [16] on strain PA14 found 165 ncRNAs in total. Here we adopted a conservative approach by manually analyzing and reporting only on those 26 novel ncRNA transcripts that were clearly expressed based on our selection criteria [H(dRNA)cut_off = 500] for our RNA-Seq datasets (Table 3); these transcripts were subsequently confirmed by RT-qPCR (NB. four additional putative ncRNAs identified from the RNA-Seq data could not be confirmed by RT-qPCR.). As controls, 5 of the 63 previously-identified ncRNAs [15, 34], (*P1*, *rsmY*, *prrF1*, *prrF2* and *prrH*) were PCR confirmed here (Table 3). 30 of the 31 ncRNAs verified through RT-qPCR were differentially expressed (Table 3, Figure S5) under one or both conditions of biofilm growth or swarming motility, which represent complex adaptive lifestyles in *Pseudomonas*.

**Table 3:**
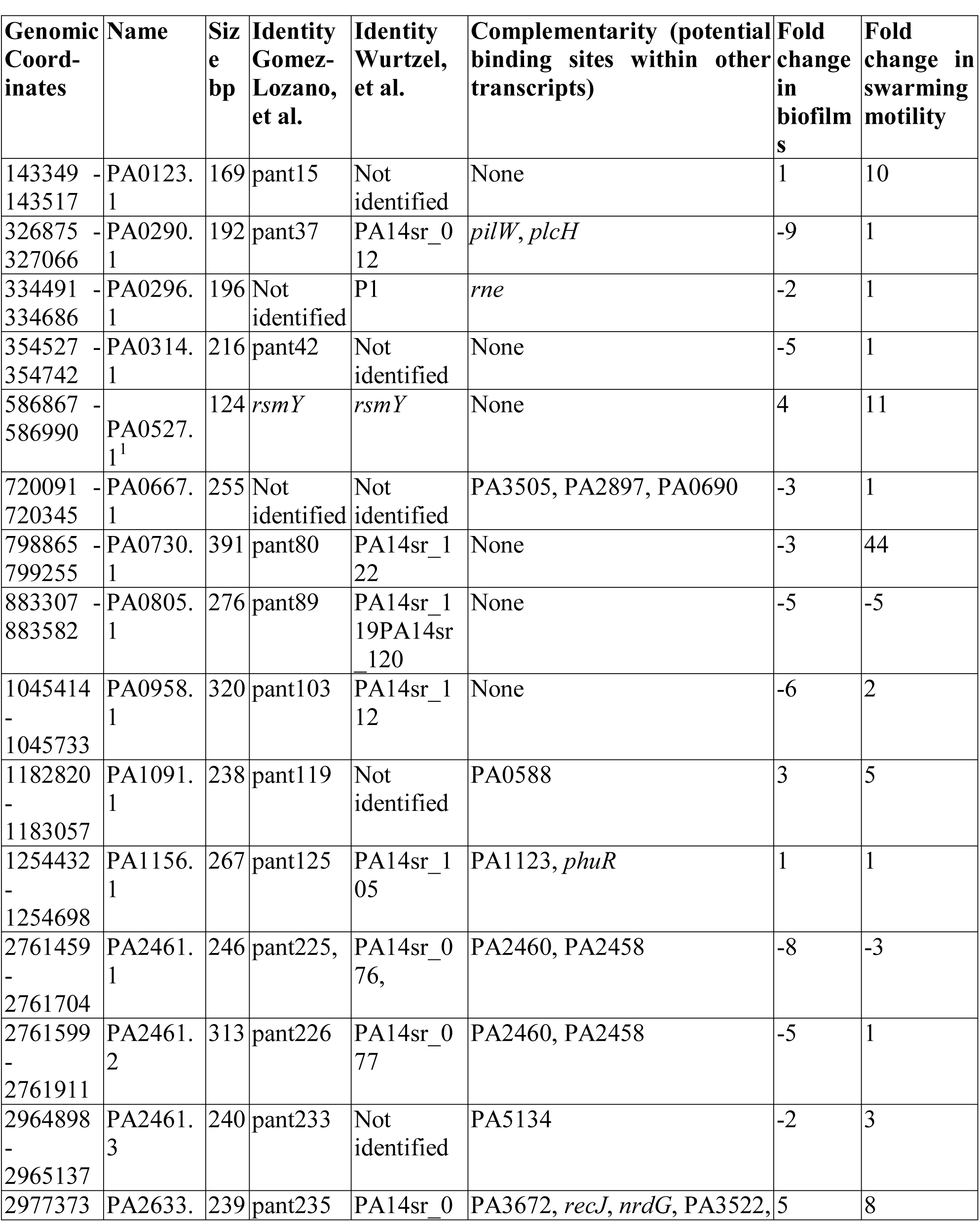
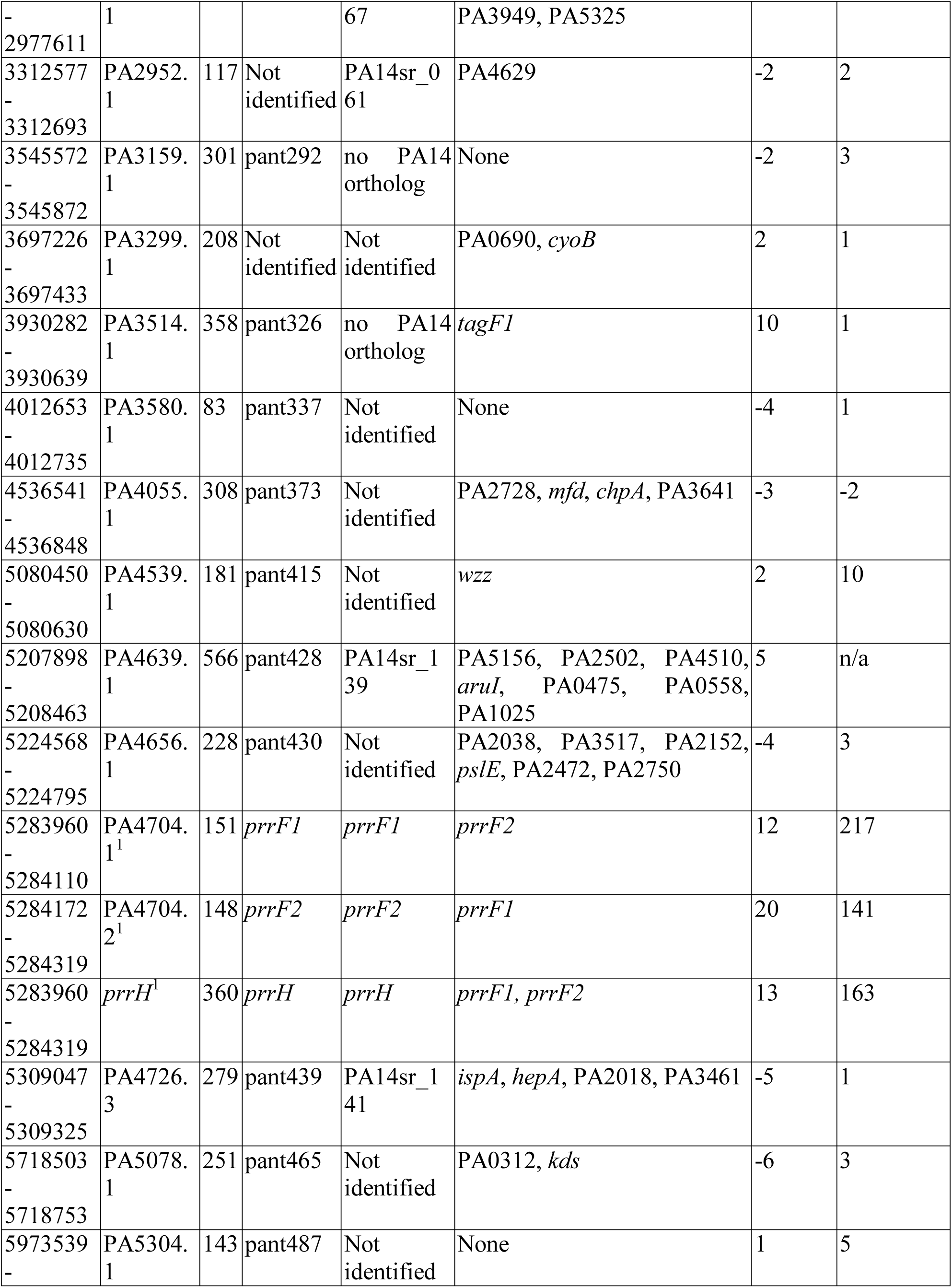
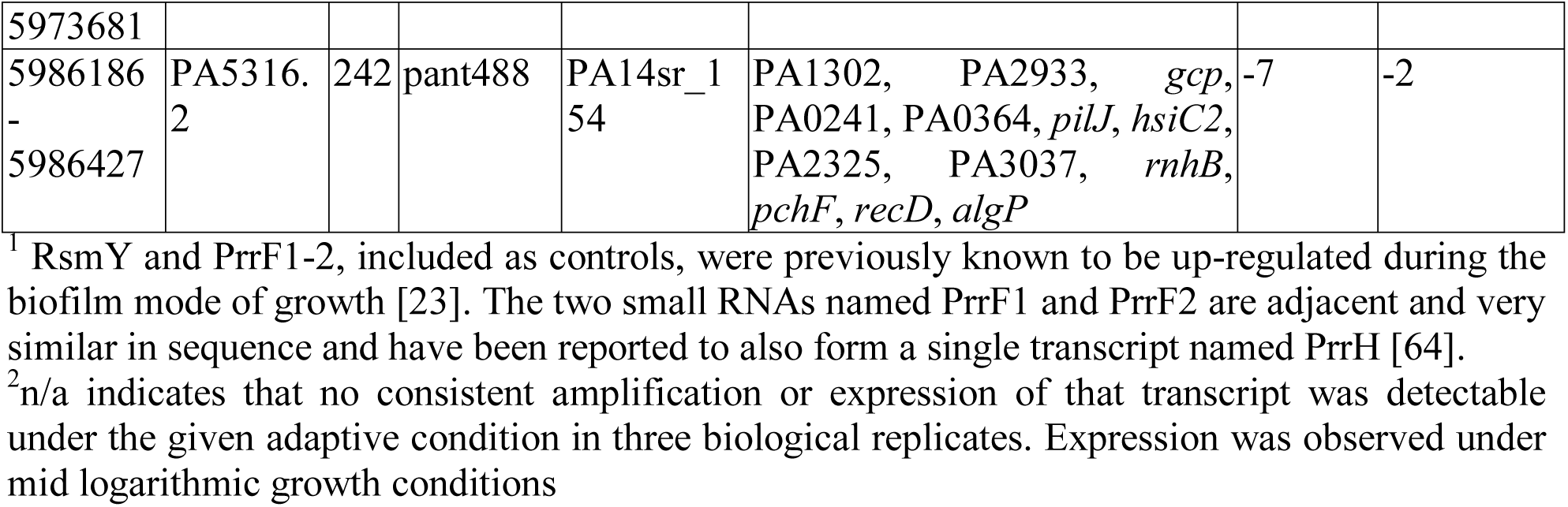
Small RNA species detected reliably by RNA-Seq and confirmed by RT-qPCR. Differential expression in biofilms and during swarming motility. Four other putative sRNAs (68836 - 69271, 707395–707685, 830970–831031, and 99801 - 100048) could not be confirmed by RT-qPCR.

## DISCUSSION

Here we introduce a novel form of sequencing that characterizes post-transcriptional processing events that we call pRNA-seq. In addition, we performed dRNA-Seq to investigate transcriptional start sites. This marriage of existing methodology with novel techniques provided new insights into the prokaryotic transcriptome and its regulation. We have provided a comprehensive map of cleavage and TSS sites for strain PAO1 grown under standard laboratory conditions (37^0^C in LB media in the logarithmic phase of growth), characterized the associated promoters, identified RNA cleavage motifs, and further characterized ncRNAs.

### >Frequent RNA cleavage events as likely intrinsic check-points for RNA degradation

The continual turnover of RNA transcripts in a bacterial cell is a highly dynamic process that has been fine tuned by evolution. Our analysis of RNA post-transcriptional processing revealed three important steps involved in the eventual destruction of RNA species. Each process has a significant ramification for understanding the regulation of RNA turnover in the proteobacteria.

Of the 240 most abundantly transcribed protein-coding genes, the transcripts from 111 (46%) were found as cleaved products in the dRNA-Seq library (Table S2, bolded rows indicate transcripts also found in the pRNA-Seq library). Therefore, there appears to be a partial correspondence between the level of expression and the level of transcript cleavage. The observation that only about half of the most abundantly transcribed genes were processed, and that this occurred at specific sites, is consistent with the hypothesis that transcript processing reflects a regulatory process that serves to control RNA decay rate for only some of these abundant RNAs [35]. Intriguingly, nearly 18% of the most abundantly transcribed genes were ribosomal proteins, many of which were apparently processed, supporting the concept that the production of the protein synthesis machinery in actively growing cells is at least partly modulated by 5’ monophosphate generating cleavage events.

Our data indicate that cleavage sites in transcripts from protein-coding genes often occur in the 5' untranslated regions of transcripts. We propose that this might be a post-transcriptional regulatory mechanism leading to modulation of translation. In addition, 1.5% of genes (19/1238) that possessed recognizable RBS produced transcripts that were cleaved such that their RBS were removed, even though the associated gene remained intact, implying a further level of translational suppression.

Remarkably, nearly all of the 500 cleavage sites from protein-coding transcripts in our data set corresponded quite closely to a sequence motif that, based on studies in *E. coli*, would be predicted to be cleaved by RNase E: [(G,A)(C,A)N(G)(G,U,A) ↓ (A,U)(C,U)N(C,A)(C,A)][36]. Our data provides evidence *in vivo* for the widespread distribution and functionality of this cleavage motif, suggesting that RNase E plays an instrumental role in regulating the processing of an unprecedented number of cellular RNAs in *Pseudomonas* and, by extrapolation, the eubacteria. *P. aeruginosa* RNase E has 64% amino acid sequence identity to that of *E. coli* and the conservation of RNase E cleavage patterns across the γ-proteobacteria appears likely given the essential function of this enzyme [3, 6, 37].

*P. aeruginosa* has 66.6% G+C in its genome, therefore the third codon positions in this organism tend to be occupied by guanine and cytosine. Our analysis indicated that cleavage sites tended to be located between the first and second codon positions in annotated protein coding genes. Thus most of the cleavage sequence motifs showed a preponderance of G and C residues in their third codon positions (the -2, +2, +5, etc. positions within the cleavage sequence motifs) (Figs. 5B and S1A). For example, the G residue at position -2, which is known to confer rapid RNase E cleavage in *E. coli* [36], was found to be conserved in 65% of both the Sharp and Tail L motifs. The notable exception to this trend was the Tail R motif, where a U residue was conserved in more than 65% of the sequences at the +2 location downstream of the cleavage site, at a 3^rd^ codon position (see Fig 5D). As this motif did not match any previously described motifs for known nucleases, the nuclease performing this cleavage, while possibly unique, is currently unknown. Sequence motifs such as Tail L, that showed RNA cleavage patterns 5’ of the cleavage site identified for the predominant Sharp cleavage site (Fig. 4C), likely resulted from a two-step cleavage process where a primary cleavage event would serve to recruit subsequent nuclease complexes that would ultimately degrade RNA in the 5’->3’ direction (consistent with a degradosome type of activity) [3].

A total of 119 peaks corresponded exactly to a TSS found in our dRNA-Seq library. As mentioned above, we believe that these are mRNAs that have been matured by a pyrophosphatase and are consequently bona fide entities within *P. aeruginosa* cell. tRNAs and tmRNA (both require 5’ monophosphates to be biologically active) were present in the dRNASeq library, but not in the TSS library. It is possible that some TSS were identified incorrectly due to incomplete enzyme digestion during library preparation. However, since we do not see any tRNAs or tmRNA in the TSS libraries, we can conclude that enzyme digestion (and elimination of transcripts possessing 5’ monophosphates) from the TSS library was successful. Therefore, we can also conclude that the overlapping sites from the dRNA and pRNA-Seq libraries exist within the cell, and that such transcripts are processed by pyrophosphatases after transcription. Triphosphates at the 5’ end of *E. coli* transcripts can be removed by the pyrophosphatase RppH; thus it seems possible that the homolog YgdP in *P. aeruginosa*(67% identity to *E. coli* RppH), may be responsible for this activity in *Pseudomonas*.

RNA cleavage resulting in RNA with a 5’ monophosphate was associated with a surprisingly diverse class of genes that had not been previously thought to be subject to such regulation and was found at notable levels in a surprisingly high number of protein-coding genes. Genes of note included PA3648/opr86 (an Omp85 homolog), which encodes the only known essential integral outer membrane protein in *P. aeruginosa*, involved in outer membrane biogenesis. This gene and many other coding genes were not known to be subject to RNA-cleavage based regulation. Notably, the impacted protein-encoding genes tended to code for essential cellular functions. Our analysis represents a starting point for more in-depth characterization of the important role of RNA cleavage in transcriptional regulation and overall cell stability.

### A comprehensive TSS profile highlighted correlated antiparallel transcription and alternative promoters

Our observation of 105 back-to-back TSS spaced by 18 bp and often containing a palindromic A/T motif at the precise center of the back-to-back TSS has at least two mechanistic explanations. First, an individual polymerase holoenzyme complex could bind to either motif and form an open-form transcriptional bubble in the palindromic region through sigma factor specific interactions. Such complexes might transiently prevent transcription in the opposite direction and/or provide a competitive mechanism for transcriptional initiation. Second, we propose an additional model where transcription is potentially initiated by an RNA polymerase dimer. In this model the spacing of the polymerase active sites on the dimer would be responsible for the 18 bp spacing observed in our data, while the palindromic A/T motif would facilitate the opening of the DNA duplex so as to allow either unit of the dimer to compete for transcriptional initiation (Fig.7). This model would therefore predict approximately equal transcriptional initiation in either direction for a fully palindromic site and biased transcriptional initiation for an asymmetric site. This second model is consistent with the finding that RNA polymerase dimers have been observed during the purification of bacterial RNA polymerase for crystallography [38].

**Figure 7:**
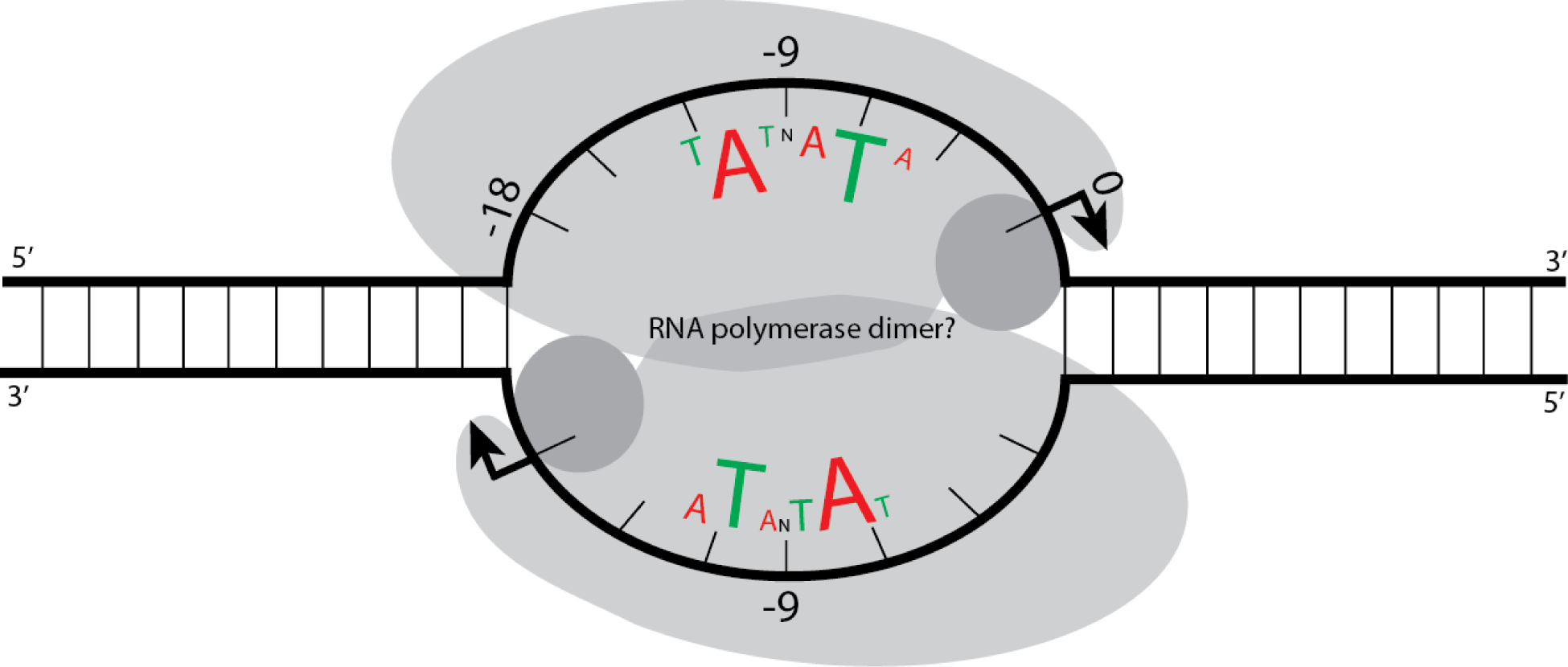
Proposed polymerase dimer binding to a palindromic antiparallel TSS. A polymerase dimer is shown bound to a region containing antiparallel transcription start sites, where transcription could take place from either or both start sites. The polymerase active sites are depicted as shaded gray circles. Within the region where the polymerase dimer is bound, only every third base is shown for simplicity. The two transcription start sites are 18 bases away from each other.

Previously, dRNA-Seq studies have been shown to be an accurate method for determining prokaryotic TSS in *Helicobacter pylori*, *Anabaena* sp. PCC7120, *Trichodesmium erythraeum* IMS101 and *Burkholderia cenocepacia* J2315 [19, 39–41]. In this study, we have effectively expanded the dataset of probable TSS in PAO1 by a factor of thirty. Our variation from published data (78% of our TSS lie within ± 2 nt of the 51 previously described strain PAO1 TSS [14] (Fig. S3)) is slightly larger than that observed by the *H. pylori* group who pioneered dRNA-Seq methodology [19] (87% of TSS within ±2 nt of published TSS). This might reflect the possibility that multiple promoters are used to transcribe the same gene, experimental errors, or variations in growth conditions in the previously published studies cf. our study (e.g., different OD_600_ at harvest) resulting in the use of different TSS. *P. aeruginosa*, with its notably larger genome and number of transcriptional regulators cf. *H. pylori*, might have a more complex transcriptome. Overall, our predicted TSS showed a mean deviation of 5.0 bases from previously published TSS. Most of this variation was due to the presence of 6 outliers that had differences from published TSS ranging from 8 to 66 nt. Interestingly 3 of the 6 outliers were involved in transcriptional regulation, and may have multiple promoters enabling different transcriptional hierarchies depending on the growth conditions.

Wurtzel et al. [16] greatly expanded the catalogue of annotated TSS in *P. aeruginosa* strain PA14 by employing a 5’ triphosphate transcript mapping strategy. Strains PA14 and PAO1 are highly similar organisms, with genomes differing by only roughly 200 genes [42]. In both the PA14 study and the present study, the bacteria were grown in identical media, and very similar TSS mapping methods were used. The number of TSS identified in PAO1 in the current study was 3,159, while Wurtzel et al. [16] identified 2,117, a difference of more than 1,000. The difference likely lies in the 25-100 fold number of total reads mapped to the non-rRNA regions in the genome in these studies (29,801,000 reads in our 5’ triphosphate library vs. 218,000 reads in Wurtzel et al.’s 37 degree 5’ triphosphate library, and 1,262,000 reads in their 28 degree 5’ triphosphate library [16]). This greater sequencing depth enabled more stringent cutoffs to be employed here than in the PA14 study (a threshold of 500 reads mapping to a single genomic location, cf. a threshold of 5 reads in PA14) [16].

In many bacterial RNA-Seq studies, it appears to be fairly common to find transcripts generated from the DNA strands opposite to those on which ORFs are located. This has been termed “antisense transcription” [19, 23]. In our study, antisense transcripts were 7% of the primary transcriptome, as predicted by mapping TSS locations. Such transcripts were found to be 12% of the transcriptome of strain PA14 [23], while in *H. pylori*, they represented 27% of the transcriptome [19]. The large difference between these organisms might reflect different cutoff thresholds used for determining what qualifies as a TSS. It could, however, reflect the biological differences that exist between *P. aeruginosa* and *H. pylori*, or reflect the fact that the *P. aeruginosa* genome is 3-fold larger than that of *H. pylori*.

A RNA-Seq study of *P. aeruginosa* PA14 focused on the changes in gene expression between cells grown in planktonic vs. biofilm conditions [23]. This group interpreted the first base at the 5’ end of RNA-Seq read pileups as the TSS. They identified a total of 3,389 putative TSS (1,054 of which were present under more than one culture condition). Their study demonstrated a high degree of reproducibility between biological replicates as well as between different culture conditions, confirming that RNA-Seq can be used to detect expression of genes encoding essential proteins and proteins involved in housekeeping functions, as well as those genes that are environment-specific. However some differences were evident between this prior study and ours. Dotsch et al. [23] reported that 75% of TSS are upstream of start codons in strain PA14 based on conventional RNA-Seq, while here we determined by dRNA-Seq that only 55% of TSS are located upstream of start codons in strain PAO1. This might be due to differences in methodology, and we note that the dRNA-Seq method demonstrated similar proportions of transcripts (49%) beginning upstream of start codons in the distantly related ε-proteobacterium *Helicobacter pylori* [19]. The use of standard RNA-Seq data would make discerning the 5’ ends of reads that form peaks in the middle of actively transcribed genes very difficult due to the cDNA fragmentation process inherent in library construction and the nucleolytic degradation that occurs rapidly with labile prokaryotic RNA. In contrast, the dRNA-Seq methodology enables the identification of the 5’ ends of transcripts regardless of where they lie within a gene, and the method is not sensitive to nucleolytic degradation that might otherwise obscure TSS signals within genes. Similarly, the fragmentation process that occurs during library construction will not obscure the 5’ ends of the cDNA molecules. Therefore, dRNA-Seq data does not share the same compounding set of interpretation issues as standard RNA-Seq data.

Under the standard growth conditions used, only 34.5% of genes in PAO1 were found to have an upstream TSS, including genes found in predicted operons. In contrast, Toledo-Arana et al. report that under all conditions studied, the firmicute *Listeria monocytogenes* transcribes at least 98% of its genes [43]. This difference likely reflects the large metabolic diversity/flexibility evident in *P. aeruginosa*[1]. Its repertoire of 24 sigma factors [9] and nearly 10% of genes involved in transcriptional regulation, is large for a bacterium, and the diverse conditions under which *P. aeruginosa* lives requires the complex interplay of genes expressed under different conditions to ensure survival and competitiveness.

### RNA-Seq data revealed novel genes and layers of transcriptional complexity

In addition to the novel layers of transcriptional complexity revealed by our dRNA-Seq analysis, this study detected other impacts on transcription. For example, data from the RNA-Seq library analysis indicated a trend towards a reduction in the level of transcription around the terminus of replication (Fig. 3). This is in concordance with Illumina DNA sequencing [31] and transposon mutagenesis studies [32] that also recovered decreased numbers of sequences and mutants respectively from this region of the genome. This phenomenon is likely related to the manner in which the genome replicates, since at any given time in an actively growing cell culture there would be more DNA present at the origin of replication than at the terminus. However, here it was demonstrated that this bias is also detectable at the transcriptional level with more transcripts evident from genes that are present in regions that are origin-proximal than terminus-proximal, likely due to a combination of gene dosage and the higher tendency for relaxation of supercoiling at the origin, which would impact gene expression. Genomic rearrangements occur more commonly in a symmetrical fashion around the terminus and origin, rather than between the terminus and origin regions [44], since such symmetrical rearrangements would conserve existing levels of gene expression by enabling genes to maintain the same distance from the origin of replication, and therefore the same copy number. In contrast it can be anticipated that there would be fitness costs to the organism if origin-proximal genes were relocated near to the terminus as supported by previous studies of detected genome rearrangements in *P. aeruginosa* [44].

Small, non-coding RNA (ncRNA, sRNA) transcripts are emerging as a major mechanism for regulating translational expression in bacteria and were also investigated here. At the time of the current study there were 140 annotated ncRNAs in PAO1, based on the *Pseudomonas* genome database [15] and the Rfam website [45]. All but 63 of these are rRNAs and tRNAs. Recent *P. aeruginosa* transcriptome investigations have sought to more thoroughly annotate ncRNAs. A recent RNA-Seq study [33] on strain PAO1 grown in LB at 37^o^C into exponential and early stationary phase suggested an additional 513 ncRNAs in PAO1. However this was based on low-stringency, automated computational methods. The existence of several of these ncRNAs was verified by Northern blot. Conversely, others [16] identify 165 novel ncRNAs in their transcriptome analysis of *P. aeruginosa* PA14. Many of the ncRNAs defined in that study employed a very low threshold for TSS prediction (minimum number of 5’ triphosphate library reads aligned to a single genome position = 5), as well as a small number of total RNA-Seq reads spanning the region of the predicted ncRNA. Therefore, these ncRNAs predicted by others should be further verified using additional sequencing data, or an additional method such as Northern blotting or PCR, to confirm their existence and expression under different conditions, as well as looking at their potential regulatory roles in the cell. Our current investigation utilized high stringency methods and confirmatory RT-PCR to define 31 reliable ncRNAs that were produced under the investigated conditions, and not all of these were identified in the abovementioned RNA-Seq studies. Intriguingly 30 of these were found to be dysregulated under the adaptive lifestyle conditions of biofilm formation and/or swarming motility. This indicates that it is important to consider different growth conditions when confirming ncRNA species and implies that translational regulation mediated through ncRNAs may be an important element in determining adaptive lifestyle changes. Certainly this is true of the ncRNA *crc* [46] and *rsmYZ* [47] and we have preliminary evidence implicating the importance of *prrF1,2* and *phrS* as critical regulatory elements in one or both of these lifestyle changes.

## CONCLUSIONS

Here, novel the pRNA-Seq methodology is reported, and shown to be a powerful tool to identify RNA transcript 5’ monophosphate cleavage/processing sites in a genome-wide manner. Cleavage sites were predominately located between the first and second codon positions within protein-coding genes. Further examination of cleavage sites has revealed that they can be classified into five distinct categories, based on their cleavage peak shape and associated sequence motifs. The cleaved transcripts occur in genes associated with specific KEGG categories, but a much wider set of categories and genes was observed than initially anticipated. We also identified a correlation between TSS that lie ∼18bp apart on opposite strands of the transcriptome. These sites are separated by a distinct motif, and transcription may be initiated here by RNA polymerase dimers. This combination of pRNA-Seq, dRNA-Seq and RNA-Seq thus provided us with a more extensive view of the transcriptome.

Due to the diversity of lifestyles and complex adaptations that *Pseudomonas* can undertake, it would be necessary to perform similar analyses, under these conditions, to those described here in order to define the transcriptional complexity that underpins diversity in this organism. Nevertheless, this study has provided a new window into a previously unappreciated level of complexity in RNA processing in a bacterial transcriptome. It involves a greater extent of transcript cleavage than previously anticipated, evidently occurring in a regulated fashion through enzymatically-controlled processes. However, this is only the start, as we expand, in the future, our understanding of the role and significance of RNA processing events in maintaining a dynamic, flexible and robust bacterial transcriptome.

## METHODS

### DNA extraction, genomic library construction and sequencing

*Pseudomonas aeruginosa* strain PAO1 was grown in Luria Broth (LB) medium at 37^0^C (to OD_600_ =∼0.7 at ∼200 rpm. DNA was extracted using a protocol modified from Cheng and Jiang [48]. Briefly, cells were collected by centrifugation at 4^0^C, washed twice with STE buffer (100 mM NaCl, 10 mM TRIS-HCl, 1 mM EDTA, pH = 8.0), then resuspended in TE buffer (pH =8.0). Cells were lysed by adding phenol and vortexing for 60 seconds. Chloroform phenol extractions were performed to extract DNA. DNA was precipitated with ethanol and sodium acetate. DNA was sheared and size fractionated on a 10% SDS-poly acrylamide gel electrophoresis (PAGE). The 190-210 bp region was excised, and DNA eluted and purified using a QIAquick purification kit (Qiagen). Libraries were constructed using the Illumina Genome Analyzer protocol and 50 bp paired-end sequence reads were obtained using an Illumina Genome Analyzer II according to the manufacturer’s instructions.

### SNP/indel analysis

Whole genome alignments were performed against the reference strain PAO1 (NC_002516) genome using 4 different tools: Bowtie [49], BWA [50], mrsFAST [51] and SSAHA2 [52]. All default parameters were used with the exception of minimum and maximum insert size specifications of 50 and 1000 for Bowtie and SSAHA2, kmer=13 and skip=2 for SSAHA2, and an edit distance of 3 for mrsFAST. After read alignments, single nucleotide polymorphisms (SNPs) were identified using Samtools [53] version 1.12 and were filtered for SNP quality scores greater than 90, read depth greater than 50 and percentage of non-reference bases greater than 90%. All heterozygous calls were removed since only a single allele is expected for haploid genomes. Most predictions overlapped across the 4 different alignments for this highly filtered set of SNPs, with the exception of BWA calling an insertion in place of 2 consecutive SNPs.

### RNA extraction

*P. aeruginosa* strain PAO1 was grown in LB medium at 37^0^C (libraries A06027, 110817_SN865 and PA0004) or 34^0^C (library PA0001), synthetic cystic fibrosis medium (SCFM) [54], and artificial sputum medium (ASM) [55] after inoculation from an LB culture grown overnight at 37^0^C at ∼200 rpm. For growth temperatures, sample cultures OD_600_ and rates of shaking, see Table 1. RNA was extracted using Qiagen’s RNA Protect Bacteria Reagent and RNeasy Midi kit (Qiagen) using the manufacturer’s protocol except that all centrifugation steps were carried out at 4^0^C. A DNase digestion step was carried out on the columns as per the manufacturer’s protocol. After elution from the columns, the RNA was further purified using Trizol (Invitrogen) following the manufacturer’s directions. rRNA depletion was performed twice using Ambion’s MICROBExpress kit. The above protocol was used for cells grown in all media with the following exceptions: A second DNase digestion was conducted for the LB 34^0^C and the SCFM samples. The OD_600_ for the samples grown in ASM media was not determined due to biofilm formation in this medium; these samples were grown for 48 h before harvesting. ASM cultures were stabilized by adding an equal volume of RNA Later (Ambion), incubated at 23^o^C for 10 minutes, then centrifuged for 30 min at 3,200 x g, 4^0^C. The pellet was resuspended in Sputasol (Oxoid) in order to break down the biofilm structure and incubated at 37^0^C for 25min with shaking. Two volumes of RNA protect were added, then the mixture was incubated at 23^°^C for 5 min and centrifuged for 30 min at 3,200 x g, 4^0^C. The supernatant was removed, and the pellet was further extracted using the RNeasy kit beginning at step #6 of the manufacturer’s protocol.

### pRNA-Seq library construction, sequencing and read mapping

The pRNA-Seq library construction process is depicted in Fig. S6. The cleavage site enriched library A06027 (LB 37^0^C) was prepared by first ligating a 17 base long adenylated DNA oligonucleotide (17.71; Table S7) onto the 3’ end of the 2x DNase treated RNA using T4 RNA ligase. A second 17 base long ribonucleotide (17.50, see Table S7) was then ligated onto the 5’ end of the RNA using T4 RNA ligase. These RNA/DNA hybrid molecules with known adapter sequences at the 3’ and 5’ ends were then reverse transcribed using 16 base oligonucleotide 16.16 (see Table S7) as a primer and Superscript III reverse transcriptase (Invitrogen). The resulting cDNA was PCR amplified using oligonucleotides 16.16 (16 bases) and 17.53 (17 bases) (Table S7) and resolved on a 6% denaturing-PAGE gel. A gel fragment containing cDNA fragments ranging in size from 600 bp - 10 kb was excised to remove any small fragments. The excised DNA was eluted and again PCR amplified using 16.16 and 17.53 to enrich for full-length cDNAs. The cDNA was then sheared and again size fractionated via SDS-PAGE. The 190-210 bp area (which corresponded to the desired library fragment size) was excised, eluted and purified with a QIAquick purification kit (Qiagen). Libraries were constructed using the Illumina Genome Analyzer protocol and paired-end 75 bp sequence reads were obtained using an Illumina Genome Analyzer II according to the manufacturer’s instructions. All reads were from the A06027 library were checked for passage of Illumina quality standards, then converted into FASTQ format. An in-house Perl script was used to trim off the 5’ ends of any reads from library A06027 that had the 17bp adapter attached from the library synthesis process. The script (provided in Supplementary Information) allowed for up to three mismatches. The trimmed reads were then aligned using Bowtie [49] as for standard RNASeq reads. Samtools [53] was then used to separate the individual reads into two strand-specific files.

### dRNA-Seq library construction, sequencing and read mapping

The RNA used to construct library SN865 (LB 37^0^C) was prepared as described previously [19] by Vertis Biotechnologie (Germany). The library was a single end Illumina library, and 50 bp strand-specific sequence reads were obtained using an Illumina Hi-Seq 2000 machine according to the manufacturer’s instructions. All reads were quality checked according to Illumina standards and then converted into FASTQ format. Reads were aligned to the PAO1 genome using Bowtie [49] except that the -X 1000 command was not used as this was a single-end library. Samtools [53] was also used to separate the reads into two strand-specific alignment files.

### RNA-Seq library construction, sequencing and read mapping

The RNA used to construct libraries PA0001 (LB 34^0^C), PA0004 (LB 37^0^C), A03674 (SCFM) and A06026 (ASM) was reverse transcribed using random hexamers and the SuperscriptTM Double Stranded cDNA synthesis kit (Invitrogen). cDNA was sheared and size fractionated using SDS-PAGE. The 190-210bp area was excised, eluted and purified with a QIAquick purification kit (Qiagen). Libraries were constructed using the Illumina Genome Analyzer protocol and paired-end 50 bp sequence reads were obtained using an Illumina Genome Analyzer II as per the manufacturer’s instructions. All reads were checked for passage of Illumina quality standards. Reads organized into FASTQ files were aligned using Bowtie [49] to the PAO1 genome using–X 1000 such that only mate pairs were reported if separated by less than 1,000bp. All other settings were the defaults. Once aligned, Samtools [53] was used to remove duplicates and select for reads that were aligned in proper pairs. The number of reads aligned to RNA genes were summarized using coverageBed (part of the bedtools software package; [56]). Reads per kb/million reads (RPKM) was calculated as a measure of expression of all genes individually under all conditions [57] using the formula: RPKM = number of mapped reads / total number of reads / gene length x 1,000,000,000

### Cutoff identification: Peak Height Threshold Determination for TSS and RNA cleavage site predictions

Please see Supplemental Information for a detailed description of how peak height cutoffs were determined for the dRNA-Seq and pRNA-Seq data.

### Prediction of TSS and cleavage sites

Once mapped, mate pairs of the 5’ monophosphate reads that were from nuclease-cleaved ends were discarded. Remaining reads from both the pRNA-Seq and dRNA-Seq libraries were trimmed of adapter sequence up to the first base pair. Genomic locations with coverage over the statistically determined cutoff (100 reads for cleavage sites/pRNA-Seq, 500 reads for TSS/dRNA-Seq as per the threshold rationale above) were marked as a peak, which represents possible cleavage sites or TSS, respectively. Regions encoding rRNA or tmRNA and their respective upstream intergenic regions were subjected to a higher cutoff threshold since the coverage in these regions was disproportionally higher. For the pRNA-Seq library, the peak height cut-off on both strands in these areas was 2,000. For the dRNA-Seq library, the cutoff on the + strand was 30,000 and on the–strand was 2,000. If multiple peaks were observed within 5 nucleotides of one another, only the location with the highest coverage was used as a predicted cleavage site. The TSS peaks were categorized according to their locations. Peaks in an intergenic region and on the same strand as the closest downstream gene = primary. Peaks within gene boundaries and on the same strand as the gene = internal. Peaks within gene boundaries and on the opposite strand from the gene = antisense. Peaks within an intergenic region = primary antisense and peaks within an area where 2 gene boundaries overlap = internal genes overlap.

### Cleavage site motif identification and function analysis

Cleavage site motifs were calculated using MEME software [58] with default parameters, except for utilizing the option to use the sense strand only, using sequence spanning from–10 to +10 bp of cleavage sites. The motif length was set incrementally from 6 bp to the full length of the sequence (21bp). The motif search did not include the reverse complement of the extracted sequence and sites within 10 bp downstream of one another were removed from the analysis, since the shared subsequence would be confusing in identifying the motif. Once an initial motif was found for all cleavage sites, sites were binned into different peak shape categories for further analysis. Peak shape refers to the number of mapped transcripts surrounding a cleavage site. A window of ± 5 bp was used to profile peak shapes. Peak shapes were subsequently categorized using k-means clustering with the R software package http://www.r-project.org/). The parameter k was estimated to be around 15 by plotting within-group variance for a number of the clusters. Therefore an initial k value of 20 was used and clusters with similar profiles were merged for subsequent analysis. An over-representation gene function analysis was calculated using hypergeometric tests based on KEGG pathway categories [28] and Holm’s test was used to correct for multiple testing.

Novel regulon member identification, RpoN binding site identification, Ribosomal binding site identification, small ncRNA identification and Functional analysis of ncRNAs by RT-PCR methods are described in Supplementary Information.

## AVAILABILITY OF DATA AND MATERIALS

All sequence data has been submitted to the NCBI short read archive. Accession numbers are as follows: [SRX156386, SRX157659, SRX157660, SRX157661, SRX157683 and SRX158075].

## AUTHORS’ CONTRIBUTIONS

EEG grew the bacteria, extracted and prepared the RNA, performed data analysis, coordinated the project and drafted the manuscript. LSC performed analysis of the cleavage site and transcription start site data. GLW performed analysis of the transcription start site data and implemented tools to visualize all sequencing data. ND helped design the pRNA-Seq protocol and prepared RNA for pRNA-Seq. RL assisted in growing the bacteria. SJHS and BKD performed sequence alignments and compared data between libraries. PKT grew bacteria, extracted RNA, conducted qPCR and performed analysis for the ncRNAs. RS conducted analysis on the transcription start sites. CS performed preliminary sequence alignments. REWH, PJU and FSLB supervised the project and edited the manuscript.

## ADDITIONAL DATA FILES

The Perl script used for 5’ end trimming on library A06027 can be viewed in Supplementary Information.

